# A cytokine receptor-targeting chimera (kineTAC) toolbox for expanding extracellular targeted protein degradation

**DOI:** 10.1101/2025.09.02.673832

**Authors:** Kaan Kumru, Zi Yao, Brandon B. Holmes, Fangzhu Zhao, Yun Zhang, Emilio Ferrara, Trenton M. Peters-Clarke, Kevin K. Leung, James A. Wells

## Abstract

Extracellular targeted protein degradation (eTPD) is as an important new modality for manipulating the extracellular proteome. However, most eTPD receptors are expressed broadly or are restricted to the liver. Cytokine receptor targeting chimeras (kineTACs) are genetically encoded bispecifics for eTPD that fuse a natural ligand like CXCL12 to an antibody, directing soluble or membrane proteins for lysosomal degradation using the widely expressed chemokine receptor CXCR7 (Pance K, Gramespacher JA., Byrnes, JR, Salangsang F., Serrano JAC, Cotton AD, Steri V, and Wells JA, *Nat. Biotechnol*. **2023**, 41, 273-281). Here, we dramatically expand the kineTAC toolbox by constructing 81 new kineTACs based on an unbiased list of cytokines, chemokines and growth factors. Remarkably, 55 of these expressed at suitable levels for analysis without any optimization. Many of these kineTACs bind receptors that have unique cell-type expression profiles, allowing for eTPD in specific cells and tissues and some were more potent than the original CXCL12-based kineTAC. We further show the internalizing capability of a kineTAC can enhance the performance of antibody drug conjugates. We believe these simple, genetically encoded tools will be useful for expanding the applications for optimized or cell type-selective eTPD.

## Introduction

Targeted protein degradation (TPD) is an exciting approach for manipulating proteins through binding and inducing their degradation.^1^ TPD provides several advantages over traditional inhibitors, including addressing therapeutically intractable targets, eliminating scaffolding functions, and overcoming resistance mutations.^2^ The TPD paradigm was first shown for intracellular proteins, which relies on ubiquitination of a target protein of interest (POI) using an E3 ligase, leading to its subsequent proteasomal degradation. However, many therapeutically important POIs reside extracellularly. Nearly three-quarters of all FDA-approved drugs target membrane or secreted proteins.^3^ Membrane and secreted proteins are most often degraded endogenously in the lysosome which has been exploited recently for extracellular TPD (eTPD) (for recent review see^4^, and prominent examples^5-15^).

A myriad of eTPD strategies have been developed to date, accomplishing degradation through the binding of glycoprotein receptors^5-6^, transmembrane E3 ligases^7-8^, Fc receptors^9–11^, and other protein or metabolite receptors.^12-13^ Most of these schemes use a bifunctional biologic consisting of a binding moiety that targets a POI such as a Fab/ScFv/V_HH_ and an additional binding moiety that recruits either a recycling receptor or a transmembrane E3 ligase to degrade targets primarily via the lysosome. Of the available receptors for eTPD^4^, with rare exception^6,14-15^ almost all are broadly expressed across many cell types. Many drugs including biologics can fail due to on-target off-disease tissue toxicity.^16,17^ Thus, access to more cell type-selective degraders could provide safer and more potent alternatives.

One of the early approaches to eTPD was the cytokine receptor targeting chimera (kineTAC), a genetically encoded bispecific that fuses chemokine CXCL12 to a POI binding antibody. The CXCL12 arm engages its recycling receptor CXCR7 to bind and degrade the POI by trafficking to the lysosome. This approach simplified the need to develop a synthetic binder for the recycling receptor, and was shown to efficiently degrade both cell surface and soluble extracellular targets.^18^ There are nearly 200 other cytokines, chemokines, and growth factors that may allow for eTPD.^19,20,21^ The receptors for these are diverse, with different tissue and cell type expression, mechanisms of internalization, and biological functions.^22^ Recognizing the untapped potential of this signaling ligand family for eTPD, we sought to develop a broad toolbox of kineTACs.

Toward this goal, we constructed 81 new kineTACs from natural signaling ligands of which 55 could be expressed and purified without optimization. We assessed their degradation potential using an internalization assay based on dye-labeled soluble vascular endothelial growth factor (VEGF) as the POI. We discovered that the kineTACs varied in rates of overall internalization, maximum internalization, and comparative effectiveness in distinct cell lines. We believe this new panel of kineTACs expand the opportunities for eTPD with greater potential for cell type specificity and efficiency and as tools for studying receptor trafficking.

## Results

### A purified protein kineTAC screen reveals a broad toolbox of degraders with diverse internalization profiles

Many cytokine receptors exhibit heterogeneous expression across tissues, especially within the immune system.^23^ As an example we analyzed the natural expression patterns of cytokine and growth factor receptors that inspired our generation of cell-type-specific kineTACs for eTPD (**Fig 1a**). We compiled immunohistochemistry data from the Human Protein Atlas^24^ for a small panel of 28 receptors known to bind signaling ligands. Expression levels were quantified across 45 normal human tissues and visualized as a heatmap (**Fig 1b**) which showed broad heterogeneity. This diversity suggested many opportunities for selective targeting.

**Fig 1.**
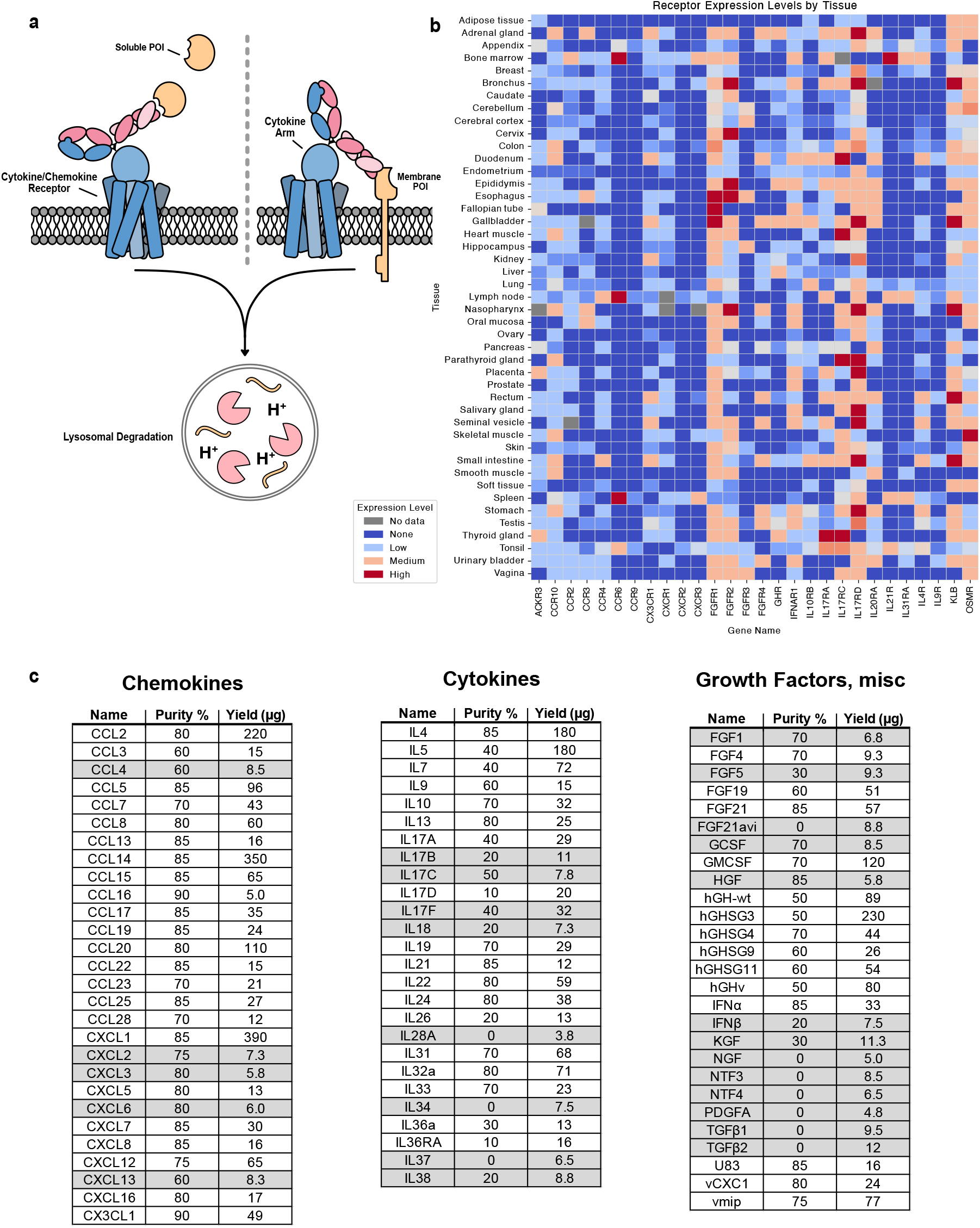
Endogenous cytokine receptor expression profiles and KineTAC library construction. **a**, The kineTAC concept depicting binding of a kineTAC to a soluble or membrane POI and to a cytokine or chemokine receptor that enables internalization and lysosomal degradation. **b**, A heatmap of some of the cytokine/chemokine receptors tested in this study and their expression patterns in human tissues illustrating their diverse distribution. Expression data from Human Protein Atlas IHC data^3^ and averaged for all available cell types in a given tissue. **c**, 81 cytokine, chemokine, and growth factor sequences were chosen and expressed as bispecific kineTACs targeting VEGF as a common POI. kineTACs shown in gray shading (26 in total) were excluded due to poor purity or insufficient yield after a single expression, leaving 55 that were tested. Purity was estimated from SDS PAGE gels; yield was estimated from A280 and molar absorptivity based on primary sequence.

We next assembled a broad and unbiased list of 81 cytokines, chemokines and growth factors as potential kineTACs (**Fig 1c**). These were cloned as knob Fc fusions and annealed to a common hole Fc-anti-VEGF (bevacizumab, beva) arm (**Fig 2a**).^25^ We chose VEGF as the model POI as it is therapeutically relevant and previously validated for degradation by the CXCL12 kineTAC.^18,26^ Importantly, having a soluble POI provided a uniform target for all the kineTACs across cell lines and permitted dose response studies. To monitor uptake of VEGF, we conjugated VEGF to a N-hydroxy succinimide pH-sensitive dye (pHrodo) which fluoresces upon trafficking to acidified intracellular compartments.

**Fig 2.**
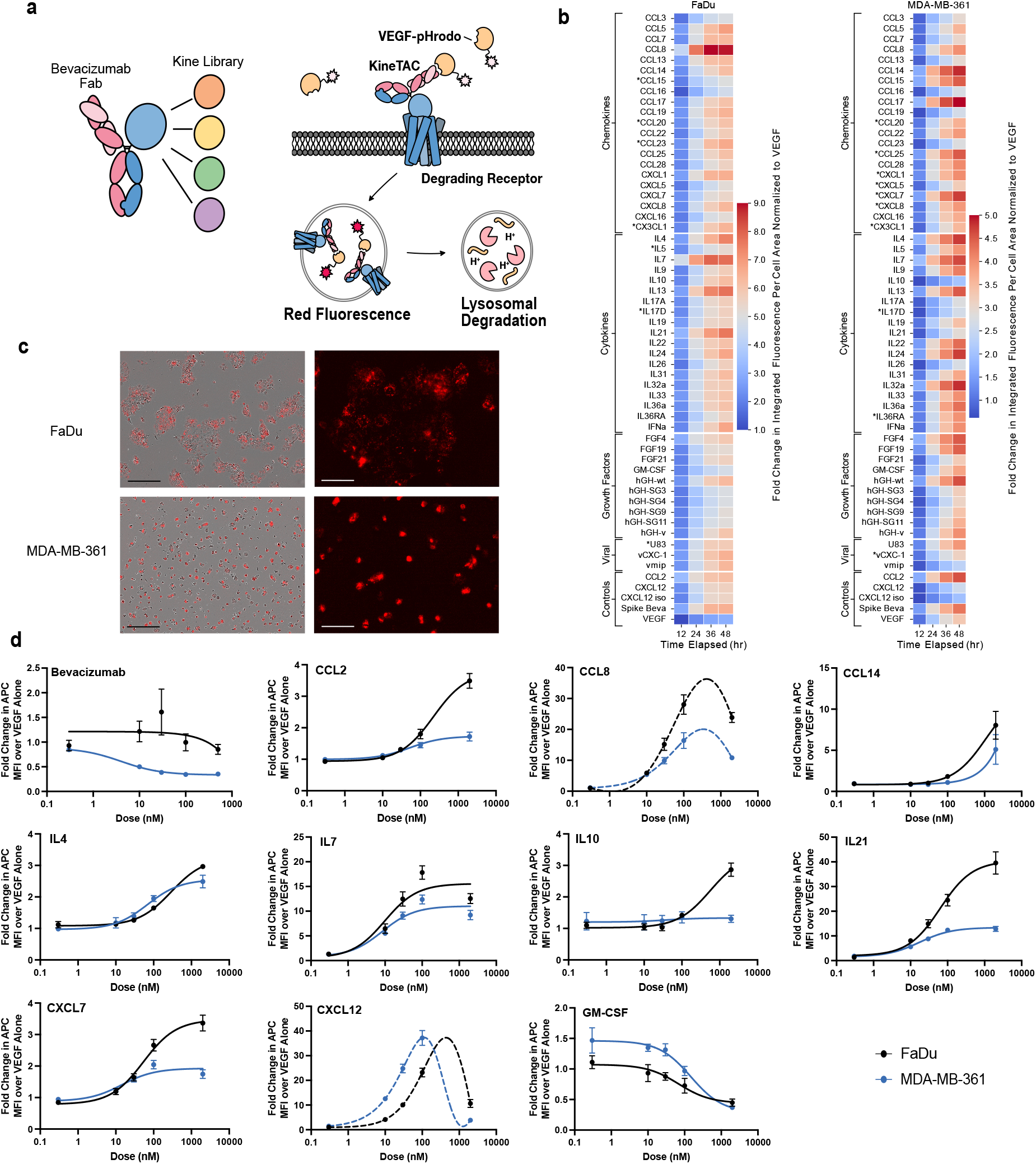
Screen unveils a range of kineTACs that can be used for eTPD. **a**, Scheme outlining the soluble POI screen that utilized fluorescently labelled VEGF as the POI. The library of 55 kineTACs (12.5nM) were expressed and purified. These were tested for internalization of pHrodo-red labeled VEGF (25nM). Internalization and degradation is evaluated by live cell imaging and detection of red fluorescence over time reflecting uptake into low pH vesicles. **b**, Screening results for two different cell lines (Fadu left, MDA-MB-361 right). Heatmap displays the fold change in integrated fluorescence intensity per cell area at given time point divided by the integrated fluorescence intensity per cell area of 25 nM VEGF-pHrodo red treated cells at 12 hrs. Data are the average of three biological replicates. KineTACs with an asterisk (*) represent no detectable mRNA transcripts for any possible receptor in the cell line (unconfirmed transcriptomics data obtained from HPA).^3^ **c**, representative images from the phase contrast/red channel only of FaDu cells (top two) and MDA-MB-361 (bottom two) at 24 hr. Red represents background subtracted red fluorescence (VEGF-pHrodo) as analyzed from the incucyte software. Black scale bar is 300 μm; White scale bar is 100 μm. **d**, Dose response curves of 12.5 nM AlexaFluor647 labeled VEGF (VEGF-647) from selected kineTACs. All kineTACs contain the displayed cytokine and a Bevacizumab half except Bevacizumab, which is the divalent IgG control. Curves are three parameter non-linear regressions, with dotted lines indicating bell-shaped curve fits.

Remarkably, of the 81 kineTACs constructed through total gene synthesis, we were able to express 55 in Expi293 HEK cells at usable levels and purities in 30 mL expression flasks (generally, ≥300 μg/L yield and ≥50% purity; **Fig 1c**). It is known that cytokines vary in their stabilities and yields of recombinant expression^27^ so we were delighted to see that so many could be expressed in this simple format without laborious optimization.

We next sought to identify cell lines expressing the receptors to test the kineTACs. Bioinformatic analysis based on nTPM from RNA seq data showed that two cell lines, FaDu (a human pharynx squamous cell carcinoma line) and MDA-MB-361 (a human metastatic breast cancer line) express receptors for all but three candidate kineTACs (CCL20, CX3CL1, IL17D are not covered) at detectable levels (defined as sum of nTPM for all possible cytokine receptors > 0, from the human protein atlas^3^), with 35 of the cytokine receptors being covered by both cell lines.^24^ We tested all kineTACs in these two cell lines over a 48-hour time course monitoring pHrodo using an Incucyte live cell imager. We observed a range of VEGF internalization patterns, with many kineTACs promoting significantly higher uptake compared to VEGF alone (**Fig. 2b-c**). Within each cell line, some kineTACs outperformed many others. For instance, in FaDu, CCL8, IL7, IL13, and IL21 induced the highest uptake at 48 hr. In MDA-MB-361, CCL17, CCL14, IL32a, and IL4 triggered the highest internalization at 48 hr. Kinetic profiles also varied depending on the kineTAC cell line combination; for instance in FaDu cells IL7 and CCL8 start to plateau around hour 36, while IL21 continues to rise up to hour 48 (**Fig. 2b**). In MDA-MB-361 cells, internalization kinetics tended to be more uniform, with a steady climb into 48 hours (**Fig. 2c**). These results confirm that many endogenous cytokine receptors can be functionally co-opted for eTPD of VEGF, with different overall kinetics and maximal internalization between cell lines.

To quantify the potency of candidate degraders, we selected eight highly internalizing kineTACs for detailed dose analysis: CCL2, CCL8, CCL14, IL4, IL7, IL10, IL21, CXCL7 compared to the isogenic Bevacizumab negative control^28^, and the CXCL12 kineTAC positive control ^18^, as well as one of the lowest performers from the screen, GM-CSF (**Fig 2d**). Most constructs showed saturable uptake, consistent with receptor-mediated endocytosis. Some showed decreased uptake at high concentrations, consistent with a hook effect seen in other bivalent binding systems likely caused by saturable monovalent binding to VEGF and the cytokine receptor.^29^ Different kineTACs exhibited variable activity across the two cell lines. For example, CCL2, CCL8, and IL21 had higher max internalization in FaDu, whereas CXCL12 was markedly more potent in MDA-MB-361. In some cases, like IL4 and IL7, the internalization was similar between the two cell lines. Overall, this screen identified a diverse and expanded library of functional kineTACs with varying internalization profiles. The differential activity observed across cell lines opens the door to selective targeting strategies based on cytokine receptor expression.

### KineTACs internalize VEGF in FaDu cells through cytokine receptor clathrin-mediated endocytosis

Endocytosis of cytokine receptors is a complex process that has implications for signaling specificity, duration, and output.^30^ Many factors affect whether and how a given receptor is endocytosed. For instance, most cytokine receptors undergo clathrin-mediated endocytosis (CME) but may also utilize caveolae-dependent internalization depending on cell type.^20^ To this end, we tested how our lead kineTACs were internalizing VEGF in FaDu cells using pathway inhibitors.

Dynasore and chloroquine reduced VEGF-pHrodo signal from CCL8-Beva in a dose-responsive manner, suggesting that dynamin and lysosomal acidification play critical roles in screen signal, respectively (**Supplemental Fig 1a**).^31,32^ We detected no reduction in signal from CCL8-Beva with up to 150x excess Fc, ruling out FcR mediated internalization (**Supplemental Fig 1b**). We also tested several endocytic pathway inhibitors: monensin (ionophore, Golgi inhibitor^33^), bafilomycin (V ATPase inhibitor^34^), cytochalasin D (actin F depolymerizer^35^), amiloride (Na^+^ channel and Na^+^/H^+^ exchange inhibitor^35^), and MG132 (proteasome inhibitor^36^) at single doses for several kineTACs (**Supplemental Fig 1c**). Notably, all tested kineTACs showed similar inhibition patterns. The reduction in internalization caused by treatment monensin and bafilomycin suggests that the route of degradation involves CME and the lysosome consistent with other eTPD systems.

To increase the sensitivity of the internalization assay, we switched to a brighter AlexaFluor647 labeled VEGF (VEGF-647) and confirmed our assay signal corresponds to internalized VEGF (**Supplemental Fig 1d-e**). Interestingly, cytochalasin D treatment increased AlexaFluor647 labeled VEGF (VEGF-647) internalization by about 2-fold, potentially due to shifting pathways from decreased phagocytosis and macropinocytosis to increased receptor-mediated endocytosis.^37^

### Evaluation of the kineTACs for membrane POI degradation

We next tested the ability of lead kineTACs to degrade membrane POIs. We first probed degradation of the key oncogenic receptor tyrosine kinase epidermal growth factor receptor (EGFR) in HeLa and FaDu cells (**Fig 3a-c** and **Supplemental Fig 1f**).^38^ A subset of nine kineTACs were converted to EGFR binders by fusing the Cetuximab (Ctx)^39^ half-IgG to a cytokine knob half-IgG. The new kineTACs ranged in their EGFR degrading potential in HeLa cells, averaging ∼40% degradation over vehicle; some were comparable or higher to that seen for our original CXCL12-Ctx kineTAC. Interestingly, the data revealed differences in degradative potential between soluble and membrane targets, which may vary even for the same cytokine. For instance, CCL8-Beva consistently increases VEGF uptake by 20-30x at 50 nM but results in a modest EGFR degradation of ∼50% in HeLa or FaDu cells (**Fig 3b, Supplemental Fig 1f**). These results confirm that kineTACs from the expanded toolbox are capable of degrading not just soluble POIs like VEGF, but also membrane targets like EGFR.

**Fig 3.**
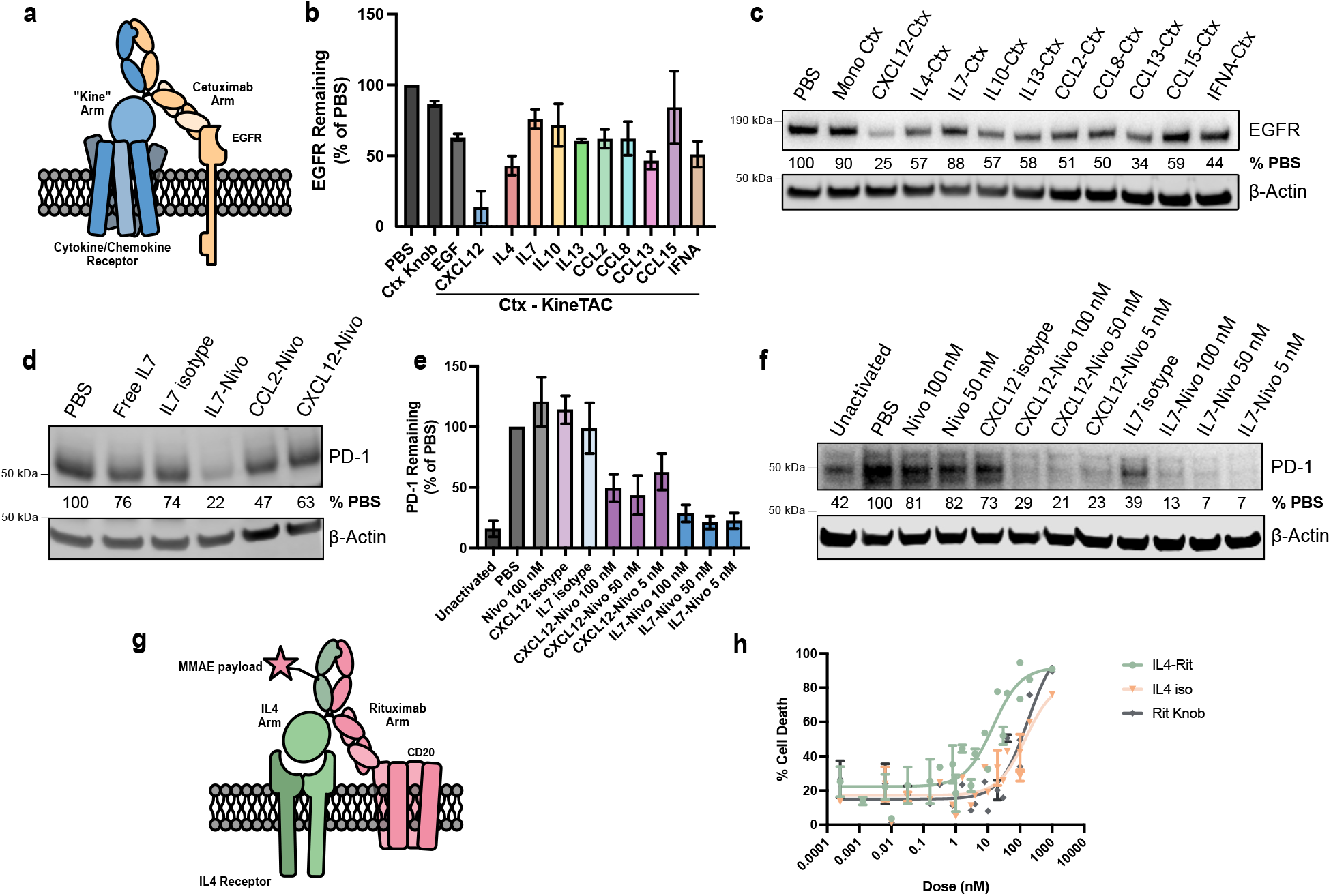
KineTAC application for eTPD of therapeutically relevant membrane proteins. **a**, scheme for EGFR degrading kineTACs includes the presented cytokine-Fc half with a common Cetuximab (Ctx)-Fc that binds EGFR. **b**, Western blot quantification of mean β-actin normalized EGFR levels as a percentage of the PBS only control after 24 hr treatment with 50 nM kineTAC or isotype Ctx single-arm control. Ctx Knob = monomeric cetuximab (Cetuximab Half IgG Hole with Fc only Knob IgG). Error bars are SEM. **c**, representative western blot that was quantified in **b**. EGFR signal is HRP luminescence and β-actin is from a LICOR secondary. The EGFR signal was normalized to β-Actin as a percentage of the PBS only normalized EGFR signal. **d**, Representative PD-1 degradation in a Jurkat PD-1 overexpression cell line after 24 hr with 50 nM of the displayed kineTAC or isotype antibody control. Nivo = nivolumab. The PD-1 signal was normalized to β-Actin as a percentage of the PBS only normalized PD-1 signal is displayed. PD-1 signal is HRP luminescence and β-actin is from a LICOR secondary. IL7 isotype = half IL7 knob IgG with Fc only hole IgG **e**, Western blot quantification of activated primary T cell mean β-actin normalized PD-1 levels as a percentage of the PBS only control after 24 hr treatment with kineTAC at displayed concentrations or isotypes at 100 nM. Nivo = Nivolumab. Error bars represent SEM. Five total donors are represented. **f**, Representative western blot of PD-1 degradation in activated donor T cells from **e. g**, KineTAC-ADC diagram. An MMAE payload is conjugated to an IL4-Rituximab kineTAC via a cleavable Val-Cit linker. Binding to both the IL4 Receptor and CD20 allow for internalization of the complex and release of the MMAE payload. **h**, cell killing of Ramos cells by the specified doses of IL4-Rituximab-MMAE or isotype-MMAE (IL4 iso = IL4 containing IgG, no rituximab. Rit Knob = Monomeric rituximab IgG, no IL4). Lines represent non-linear three parameter agonist vs. response curves. Data are from three biological replicates and error bars represent SEM.

### Both antagonist and agonist kineTACs can degrade membrane POIs

For a given kineTAC and application, modulating a cytokine or growth factor pathway may either complement or counteract the biological effect of degrading a POI. We had previously found that there was little difference in degradation efficiency for CXCL12-based kineTACs when using an agonist or antagonist of CXCL12, suggesting signaling was not critical for degradation. We further tested this concept by the addition of an agonist IL36α and the natural antagonist IL36RA in our screen (**Fig. 2b-c**). Both showed comparable internalization of VEGF-pHrodo, consistent with receptor-mediated trafficking occurring independently of downstream signaling. Similarly, a panel of hGH variants^40^ with altered receptor affinity did not show a clear relationship between binding strength and uptake in our screen (**Fig. 2b-c**), implying that other factors, such as receptor recycling or endocytic sorting, may dominate in these systems.

We further tested if degradation of EGFR can occur independently of signaling with kineTACs in a matched endogenous agonist/antagonist pair: IL1 and IL1RA (**Supplemental Fig 1g**). IL1 and IL1RA both bind the IL1 receptor, but IL1 is proinflammatory, inducing lymphocyte activation and local tissue destruction.^41^ IL1RA binds the same receptor but does not trigger activation, acting as a natural inhibitor. Both the agonist and antagonists reduced cell EGFR levels (IL1: 65% of PBS, IL1RA: 45% of PBS) (**Supplemental Fig 1h-i**). We also saw a trend toward EGFR degradation using a GMCSF antagonist. IL1-Ctx and isotype strongly upregulated phospho NF-κB after 15 minutes, whereas IL1RA-Ctx and isotype did not. These observations highlight the flexibility of the kineTAC toolbox: both receptor agonists and antagonists can drive eTPD and one can decide if signaling through the cytokine is beneficial along with degradation.

### Leveraging IL7 and IL4 kineTACs for specific, therapeutically relevant eTPD

To test the potential for tissue-specific degraders, we selected IL4 and IL7 as representative cytokines with known distinct receptor expression profiles and promising VEGF internalization for further validation against clinically relevant membrane POIs.^42,43^ IL7 has been used therapeutically to stimulate widespread T cell proliferation,^44^ and its receptor IL7R is tissue restricted to T lymphocytes^24^, so we wondered if the potential synergy between increased T cell proliferation could pair nicely with immune checkpoint downregulation. Towards this goal, we tested whether PD-1 could be effectively degraded by an IL7 kineTAC based on the PD-1 binder Nivolumab (Nivo) (**Supplemental Fig 2a**).^45,46^ Starting with a Jurkat PD-1 (Jurkat^PD-1^) over expression model, IL7-Nivo reduced total PD-1 to 36% (**Fig 3d, Supplemental Fig 2b**). In this engineered system, the difference between IL7-Nivo and isotype was enough to further motivate us to analyze the kineTAC in a primary T cell model. After activating donor CD3^+^ cells with αCD3 and αCD28 to upregulate PD-1 expression, 5nM IL7-Nivo treatment degraded PD-1 to ∼20% of vehicle after 24hr, which outperformed CXCL12-Nivo which had ∼60% remaining after 24 hr (**Fig 3e-f**). Additionally, we did not detect any decrease in IL7R levels suggesting minimal degradation of the cytokine receptor (**Supplemental Fig 2c**). These results suggest that IL7-Nivo kineTAC could be a viable selective checkpoint degrader in T cells, leveraging IL7R as both a delivery, specificity, and in principle an activation handle.

In parallel, we focused on IL4, a potent pleiotropic cytokine that has roles in stimulating T_H_2 cells and inducing class switching to IgE in B cells.^47^ IL4Rα expression is tightly controlled and is particularly high on B lymphocytes. Thus, we tested if an IL4 kineTAC could be useful for targeting B cells for eTPD. CD20 is a highly specific B cell antigen and has been used for targeting B cells for elimination by antibody-dependent cellular cytotoxicity (ADCC) in the treatment of diffuse large B-cell lymphoma (DLBCL).^48^ IL4Rα and CD20 are both detected at all stages in the patients with DLBCL, and anti-IL4Rα-CD20 bispecifics have shown some efficacy.^49^ There is also clinical interest in producing ADCs for CD20 to enhance potency beyond ADCC alone.^50^

We have recently shown that coupling the binding of low-density lipoprotein receptor (LDLR) to a POI can dramatically enhance the potency of an ADC through increased internalization of the cytotoxic payload in what we call a degrader drug conjugate (DDC).^13^ Thus, we hypothesized that an IL4 αCD20 kineTAC could trigger cell-specific enhancement in ADC internalization efficacy (**Fig 3g**). We constructed the IL4 kineTAC with Rituximab (Rit) to target CD20 (**Supplemental Fig 2d**) and observed a 56% reduction in CD20 after 24 hours in Ramos cells, a B cell line (**Supplemental Fig 2e-f**). This suggested to us that the IL4 kineTAC was internalizing as anticipated. We observed an increase in CD20 triggered by IL4 isotype (containing no CD20 binding arm) which is consistent with literature reports of IL4 biology.^51^ We next conjugated the kineTAC and single arm isotype controls with a cathepsin cleavable N-hydroxy succinimide ester-monomethyl auristatin E (NHS-MMAE) to a drug antibody ratio (DAR) of approximately 1.0 (**Supplemental Fig 3a-f**) and measured level of cell killing. Indeed, we saw a 12-fold increase in potency for triggering cell death upon treatment with a IL4-Rituximab DDC in Ramos cells compared to both isotype ADC controls (**Fig 3h**). These findings provide an example of how kineTACs can enable cell selective degradation and payload delivery with therapeutic relevance in B and T cells.

### Cell-selective degradation of soluble VEGF in complex cellular mixtures

The diverse cytokine receptor expression landscape presents opportunities for cell-selective eTPD targeting. To demonstrate this, we deployed the IL4 and IL7 kineTACs against cell lines that endogenously express reciprocal levels of their respective receptors: IL4Rα^high^ IL7R^low^ Daudi cells and IL4Rα^low^ IL7R^high^ SiHa cells (**Fig 4a, c**). Strikingly, IL4-Beva preferentially directed VEGF-647 internalization in Daudi cells, whereas IL7-Beva achieved significantly higher uptake in SiHa cells (**Fig 4b, d**). This selective uptake mirrored the relative kineTACs surface binding patterns and underlying receptor expression (**Supplemental Fig 4e-j**), confirming the ability to enable cell-specific uptake.

**Fig 4.**
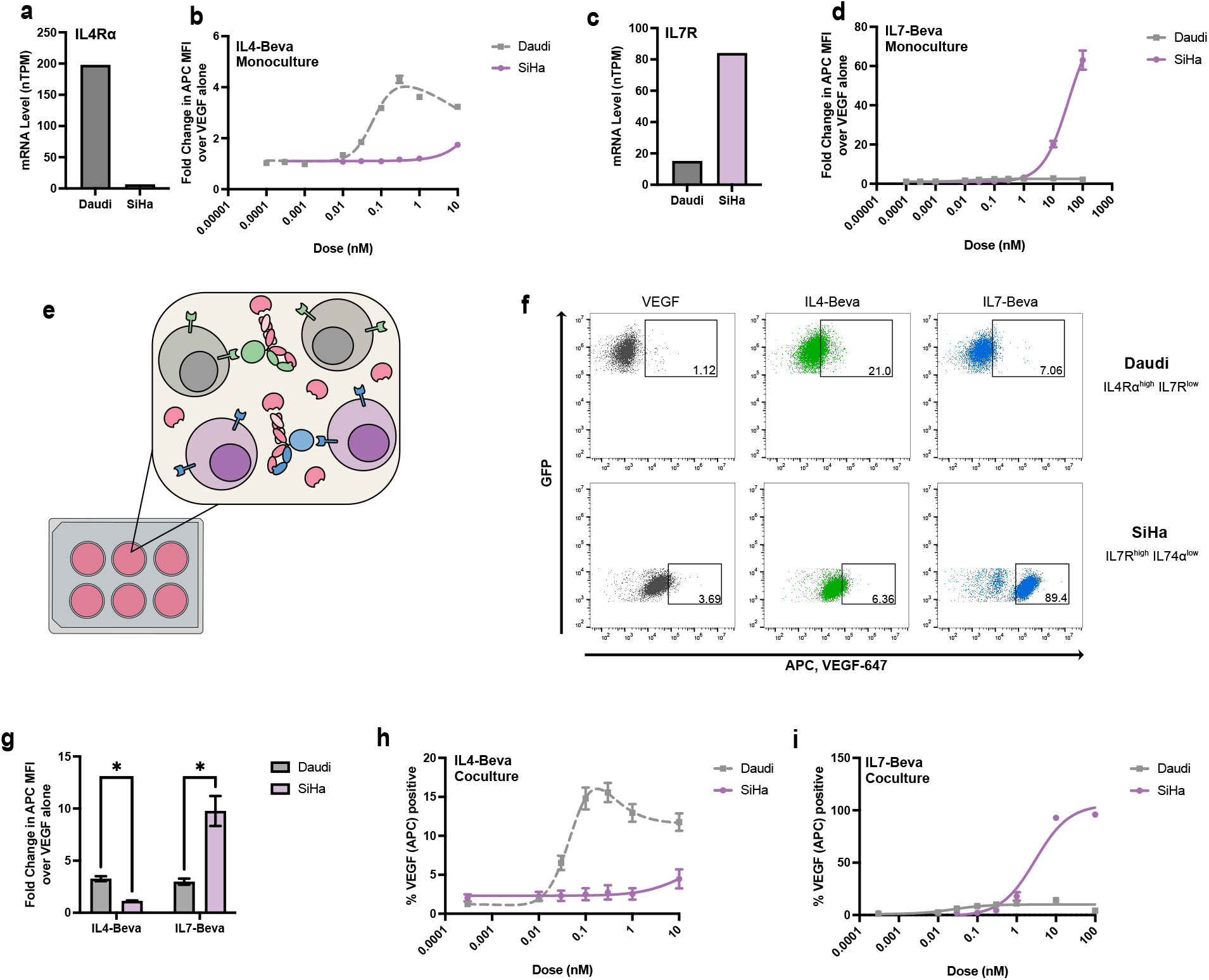
IL4 and IL7 kineTACs enable cell type-specific VEGF internalization in cell lines. **a**, mRNA normalized transcripts per million (TPM) of IL4Rα from Human Protein Atlas for Daudi and SiHa cell lines. **b**, 24 hr VEGF-647 internalization at 25 nM as a dose response of IL4-Bevacizumab in Daudi and SiHa cell lines. Curves are three parameter non-linear regressions, with dotted lines indicating bell-shaped curve fits. Means ± SEM of three biological replicates are shown. **c**, mRNA normalized TPM of IL7R from Human protein atlas for Daudi and SiHa cell lines. **b**, 24 hr VEGF-647 internalization at 25 nM as a dose response of IL7-Bevacizumab in Daudi and SiHa cell lines. Curves are best fits for three parameter non-linear regressions. Mean values ± SEM are from three biological replicates. **e**, Diagram of coculture experiments. Daudi cells (gray) and SiHa cells (purple) are co-incubated with VEGF-647 and either IL4-Bevacizumab kineTAC (green) or IL7-Bevacizumab kineTAC (blue). Cell-type specific receptor expression allows for cell-specific internalization of VEGF using the respective kineTAC. **f**, Representative flow cytometry data from coculture experiment. Daudi and SiHa cells in equal amounts were incubated for 24 hrs with 25 nM VEGF-647, and 0.3 nM IL4-Bevacizumab or 10 nM IL7-Bevacizumab. Gates show thresholds for VEGF positivity, defined as approximately 1% of total cells for VEGF-647 only. Top row: FITC^+^ Daudi^GFP^. Bottom row: FITC^-^ SiHa. Percentages in gate are displayed. **g**, Fold change in VEGF-647 median fluorescence intensity in the coculture Daudi and SiHa experiment from **f**. Mean fold change over 25 nM VEGF-647 alone ± SEM is presented. Asterisk represents a discovery (q<1%) using the False Discovery Rate to correct for multiple comparisons. **h**, Coculture VEGF-647 percent positivity (same gate as in **f**) as a dose response of IL4-Bevacizumab. Mean values and ± SEM are from three biological replicates. Curves are three parameter non-linear regressions, with dotted lines indicating bell-shaped curve fits. **i**, same as in **h**, but for IL7-Bevacizumab.

Interestingly, CXCL12 bound the surface of both cell lines (**Supplemental Fig 4g**,**j**) but triggered more internalization in SiHa cells (**Supplemental Fig 4c**). This difference correlates with receptor expression levels: in Daudi cells mRNA levels of CXCR4 are high and CXCR7 levels are low, while SiHa displays the opposite pattern (**Supplemental Fig 4a**). It is known that CXCR4 is the signaling receptor for CXCL12 and gets degraded in the process whereas CXCR7 is a recycling receptor that clears CXCL12.^52^ Because cytokines can often bind multiple receptors, we anticipate that the tissue specificity for a certain kineTAC depends not only on the exact expression of each cytokine receptor but relative internalization and/or recycling efficiency of each.

To further test the specificity of our IL4 and IL7 kineTACs, we moved to a coculture model containing both Daudi and SiHa cells (**Fig 4e**). As expected, we found that VEGF-647 was efficiently internalized into Daudi using IL4-Beva kineTAC and into SiHa using IL7-Beva kineTAC at approximate EC50 doses in coculture (**Fig 4f-g**). The dose responses for each were also largely the same as in monoculture (**Fig 4 h-i**), with somewhat lower absolute internalization but very similar dose response behaviors. In opposition to the monoculture data, CXCL12 in coculture led to similar VEGF uptake between the two cell lines that contain one or both of its receptors (**Supplemental Fig 4d)**. Taken together, these data demonstrate that kineTACs may indeed be used for cell type-specific degradation by leveraging the differential expression of cytokine receptors.

### KineTACs direct cell type-specific degradation in human PBMCs

Given the success of specific degradation in cell lines and cocultures, we sought to expand our cell type-specific eTPD panel by targeting cytokine receptor expression patterns in primary human cells (**Fig 5a**). Inspired by the restricted IL4Rα expression in B cells and IL7R expression in T cells within PBMCs (**Fig 5b-d**), we tested if kineTACs could be used to selectively direct degradation in these cell types. We isolated primary CD19+ B cells and CD3+ T cells from two donors and assayed the internalization of VEGF-647 using IL4-Beva and IL7-Beva kineTACs (**Fig 5e-f**). We found a significant increase in VEGF internalization in B cells only using IL4-Beva (mean 1.7x over VEGF alone). In contrast, donor T cells only showed an increase in VEGF-647 internalization when treated with the IL7 kineTAC (mean 12x over VEGF alone). The fold change increase with IL7-Beva was greater than that of IL4-Beva, consistent with broader expression of IL7R and higher potency of IL7-Beva as seen in the earlier dose responses (**Fig 2d**). There was some variation between the two donors in both experiments likely due to natural variation in the expression of IL4Rα and IL7R.

**Fig 5.**
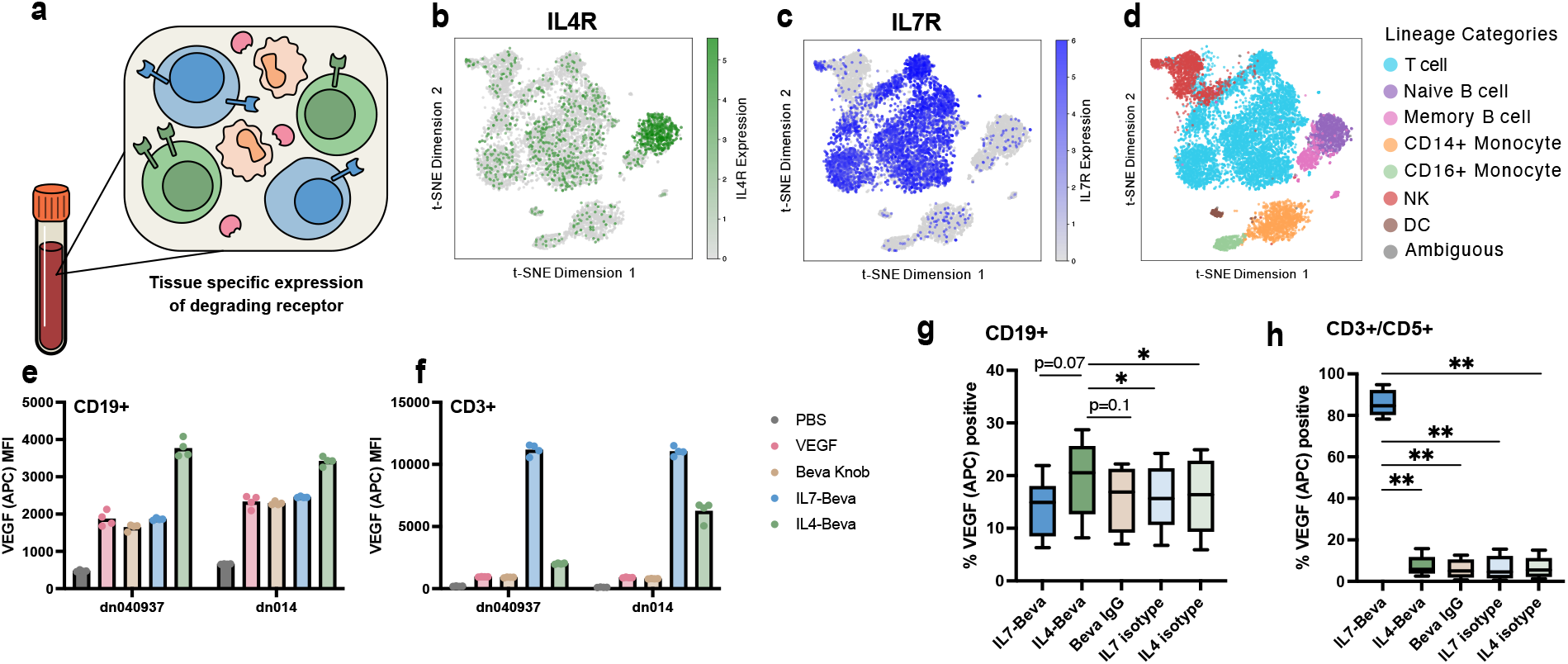
IL4 and IL7 kineTACs enable cell type specific VEGF internalization in primary lymphocytes and PBMCs. **a**, Diagram depicting tissue-specific expression of potential kineTAC receptors. **b**, tSNE analysis of IL4Rα expression from scRNA seq of two PBMC donors, obtained from the Immune Cell Atlas.^66^ Notice the prominent naïve B cell cluster of high IL4Rα expressors (green). **c**, tSNE analysis of IL7R expression from the same dataset. Notice primarily T cell restricted expression of IL7R (blue). **d**, Immune Cell Atlas cluster annotations for primary cell lineages, colored by lineage as in legend to the right. **e**, Raw VEGF-647 median fluorescence intensities across two donors for isolated CD19+ cells from four technical replicates. Cells were incubated with 25 nM VEGF-647 and 10 nM of indicated kineTAC or isotype. **f**, same as **e**, but using CD3+ T-cells. **g-h**, VEGF-647 with PBMC components. 10 nM of the indicated kineTAC or isotype was added to 25 nM VEGF-647 along with 200k cells/well PBMCs and allowed to internalize for 24 hrs. Percent VEGF positive was defined by the no VEGF-647 condition (see **Supplemental Fig 6**). Analyzed by repeated measures donor-paired one-way ANOVA, with p values for Dunnett’s correction for multiple comparisons to the kineTAC of interest (IL4-Beva for B cells, IL7-Beva for T cells) (* = p<0.05, ** = p<0.01). **g** Analysis of B cell gate and **h** analysis of T cell gate.

The promising selectivity observed in isolated B and T cells spurred us to escalate the challenge to the more complex cellular milieu of a PBMC model. We tested if kineTACs could precisely target VEGF-647 uptake to B and T lymphocytes, distinguishing them from other cell populations such as NK cells, monocytes, and dendritic cells of PBMCs (**Fig 5g-h**). Across four donors, IL7-Beva dramatically increased the number of T cells that took up VEGF-647 over Beva IgG and VEGF-647 (mean 85.4%), significantly higher than any other treatment. IL4-Beva significantly increased the number of B cells that took up VEGF-647 (mean 19.4%), of modest significance above all other treatments. The effect size of IL4 was smaller, likely reflecting the lower expression level of IL4Rα on B cells relative to IL7R on T cells that we saw in both cell lines and isolated primary cells. Gratifyingly, we also observed T cell specific surface staining at 4°C with both IL7-Beva and IL7-isotype and B cell specific surface staining with IL4-Beva and IL4-isotype, confirming that cell specificity of the kineTACs is driven by cytokine to cytokine receptor binding (**Supplemental Fig 5a-b**). Additionally, we observed a reduction in IL7R staining in T cells after treating with IL7-Beva and isotype and a modest reduction in IL4Rα staining in B cells after treating with IL4-Beva and isotype at 4°C, further supporting that these receptors are responsible for internalization as kineTAC binding blocks staining antibody (**Supplemental Fig 5c-d**). These data suggest the kineTACs can be used for tissue specific eTPD.

## Discussion

eTPD has emerged as an exciting new methodology with many uses as tools for biologists and potential therapeutics. However, the space has relied on a relatively small number of degrading receptors, most of which have broad tissue distributions that preclude cell type-specificity. In this work, we begin to address this limitation by developing and validating a large library of kineTACs, strategically designed to harness the natural distribution of cytokine receptors with greater cell selectivity. The advantages of kineTACs are that (i) they can be applied for both soluble and membrane associated targets as they rely on binding and internalizing a cytokine receptor with no change to the intracellular domain of the POI; (ii) they are entirely genetically encoded and do not require bioconjugation; (iii) they utilize natural cytokines obviating the need to generate a synthetic binder or foreign protein making them more accessible; and (iv) the cytokine is a single chain molecule and avoids issues of light-chain scrambling that challenge bi-specific IgGs. Since only a couple of kineTACs were originally described,^18^ we were enthusiastic to expand this modality to other potentially cell type specific receptors.

Cytokines and growth factors can pose significant challenges in expression without optimization^27^ so it is with some trepidation that we embarked to expand the kineTAC platform. Thus, we constructed a large panel of 81 kineTACs at the gene level. We were encouraged that 55 expressed at levels and purity that were suitable for testing without any optimization (**Fig 1c**).

To try to better understand the variation in kineTAC expression we examined the intrinsic biophysical parameters of the cytokine such as calculated isoelectric point, molecular weight, flexibility,^53^ average of hydropathy,^54^ and instability index^55^ (**Supplemental Fig 7c-f**). Unfortunately, there were not striking correlations. However, considering family level data we found that expressible cytokine kineTACs trended lower in molecular weight, expressible growth factor kineTACs had a slightly lower pIs, and expressible chemokine kineTACs had a lower instability index. There was also considerable variation in replicate expression experiments that contribute to noise. Nonetheless, we believe that bespoke optimization campaigns may increase expression levels for kineTACs. For example, in later expression tests we obtained up to 50x improvement in yields for some kineTACs by switching from the classic IL2 signal sequence in the pFUSE vector to a H5 signal sequence^56^ in a pCDNA3.4 vector, and employing an optimized CMV3.4 promoter, optimized intron, polyA signal, and improving codon bias. Thus, vector choice, promoter, and signal sequence optimization would be good to test in bespoke optimization campaigns.

The next challenge was to screen the 55 kineTACs for their degradation potential. This required finding cell lines that expressed the complementary cytokine receptors. We were able to identify two cell lines, FaDu and MDA-MB-361, that in aggregate covered 52 of these kineTACs and had 35 in common which allowed cell-line specific comparisons to be made. The screen employed a uniform VEGF-pHrodo assay as a generic soluble antigen that provided a useful measure for internalization of a soluble ligand within acidified endosomes. In most cases, kineTACs that had no cognate receptor nTPMs from HPA data^3^ in a given cell line did not perform well in the screen (**Fig 2b**). One interesting exception is CXCL7 in MDA-MB-361, which internalized surprisingly well given a lack of detectable CXCR2. CXCL7 could be internalizing through another cytokine receptor in this case, or there could be a small amount of CXCR1 or CXCR2^57^ expressed on these cells but not detected in the RNA seq data. For the most kineTACs of interest it is important to conduct full dose responses in desired cell lines. Indeed, dose responses in FaDu and MDA-MB-361 on freshly expressed kineTACs revealed higher maximum internalization in FaDu (**Fig 2d**).

Not only were these kineTACs useful for degrading the soluble VEGF, but as expected, could also degrade membrane proteins like EGFR, PD-L1 and CD20 (**Fig 3, Supplemental Fig 2**). Our inhibitor studies suggested degradation utilized the endo-lysosomal system as seen previously^18^ (**Supplemental Fig 1 a-c**). The degradation efficiency varied among these targets and between cell lines. Previous studies have shown variation can be driven by POI/degrader ratio and their absolute abundance. ^14,18^ Interestingly, we did not observe any significant correlation between receptor abundance and VEGF internalization from our screen (**Supplemental Fig 7**). The variation could also come from differences in recycling capabilities for the degrading receptors and the specific way the each bi-specific engages both POI and cytokine receptor. These are properties that are worthy of further study. We also observed that the degradation potential for the membrane protein EGFR was less than for soluble VEGF internalization which has been observed before in other eTPD platforms.^4^ We speculate that the additional conformational requirement of binding two receptors, the relative abundance of both the POI and cytokine receptor on the cell surface, and unique internalization profiles of the POIs themselves could contribute to the greater difficulty for degrading membrane vs soluble targets. The dramatically increased number of new kineTACs may help to study these fundamental trafficking parameters while providing additional optionality for eTPD of desired POIs.

Cytokines are potent signaling ligands that drive distinct pro-or anti-inflammatory pathways through cell type specific interactions. Thus, it is possible that kineTACs may complement or hinder the biological purpose of POI degradation, presenting the potential for synergy or an undesirable side effect. Here, we studied if we could decouple binding and signaling by comparing agonist/antagonist kineTAC pairs. We found little difference in degradation efficiency between the matched agonist/antagonist in screen data and in later follow up with IL1 and IL1RA, despite IL1 triggering a ∼30x increase in NF-κB phosphorylation (**Fig 2b, Supplemental Fig 1 h-k**). These data suggest that binding but not receptor activation is the critical feature in driving degradation. This more extensive study aligns with our previous report that both agonist and antagonist CXCL12 variants degraded PD-L1 with equal efficiency.^18^ It is also supported by several reports of receptor agonists and antagonists triggering cytokine or chemokine receptor internalization.^57,58^ Thus, we anticipate that other kineTACs can be used as either agonist or antagonists to effect eTPD depending on their application.

Receptor internalization has been shown to be a key requirement for an effective ADC.^60^ Recently, we have shown that a degrader based on a potent recycling receptor such as the LDLR can enhance the cell killing for an ADC, called a degrader drug conjugate (DDC).^13^ Here, we show this hybrid modality can also be applied using kineTACs. We found that an IL4-Rituximab-MMAE DDC that targets CD20 on B-cells could enhance the activity the monomeric parent Rituximab-MMAE isotype by 10-fold (**Fig 3g-h**). This suggests that kineTACs can enhance the potency of ADCs for receptors that poorly internalize.

A number of promising oncology targets such as CD47^61^, EPCAM^62^, and CD33^63^ suffer dose-limiting on-target off-tumor toxicities. Cell and tissue selective degradation is an important goal for the eTPD field that could help address on-target off tissue toxicity issue. We tested two candidate kineTACs for cell selective degradation based on IL7 and IL4. IL7 has been used clinically^44^ with a T-lymphoproliferative effect, potentially synergizing with checkpoint blockade in cancer. We confirmed the cell specific degradation of PD-1 in a Jurkat^PD-1^ model and in primary T-cells across 5 donors. The IL7-Nivolumab kineTAC out-performed CXCL12-Nivolumab in monoculture PD-1 degradation, and due to IL7’s T cell selectivity, we speculate that this difference would be enhanced in more complicated models where CXCL12 may be soaked up by any CXCR7 containing cell. In addition to the potentially cell selective killing by IL4-Rituximab-MMAE, we also found cell-selective internalization of the soluble POI, VEGF, could be directed by IL4-Beva into B cell models and IL7-Beva into T cell models, even in coculture (**Fig 4**). We further demonstrated the cell-type specificity of IL4-Beva and IL7-Beva kineTACs in primary PBMCs, triggering internalization of VEGF only in the desired B or T cells, respectively (**Fig 5**). We expect that many other yet unexplored kineTACs would behave similarly, allowing for degradation specificity following the natural expression patterns of cytokine receptors.

We believe our expanded library of kineTACs, which leverage the natural distribution of cytokine receptors, will serve as valuable tools for the greater eTPD community. These genetically encoded bi-specifics are relatively easy to construct using the natural cytokine sequences and antibodies in public data-bases such as SAbDab^64^. Effective degradation depends on the co-expression of both the degrading receptor and the target POI. This layered requirement creates a built-in logic gate for tissue selectivity that could be further strengthened through protein engineering. In particular, tuning linker architecture and binding affinities between components may further enhance selectivity by biasing degradation toward cells with optimal cytokine receptor and POI levels. We anticipate that such strategies will enable future generations of kineTACs to deliver more efficient, specific, and safer extracellular degraders across diverse tissue contexts and diseases.

## Acknowledgements

We thank Dr. Jason Gestwicki and Dr. Mark Von Zastrow for their insights, and the Wells Lab broadly for helpful discussions and expertise. We also thank Madison Seto for providing Fc-biotin for FcR competition experiments. We are grateful to generous support from NIH-1R01CA248323-01(J.A.W), NIH-R35GM122451 (J.A.W.), and the Hind Professorship in Pharmaceutical Sciences (J.A.W). K.K. is supported by an NSF Graduate Research Fellowship. Z.Y. is supported by a National Institute of General Medical Sciences F32 Postdoctoral Fellowship. B.B.H. is supported by NIH 1K08NS133290-01. F.Z. is supported by an A.P. Giannini Postdoctoral Fellowship. E.F. is supported by the NIH Ruth L. Kirschstein National Research Service Award (NRSA) Institutional Research Training Grant (T32) under award number T32GM139794. T.M.P.-C. is supported by the National Cancer Institute of the National Institutes of Health under Award Number F32CA298768.

## Author Contributions

K.K. and J.A.W. conceived and designed the study. K.K. performed the kineTAC screen and initial degradation experiments. K.K, Y.Z., F.Z., Z.Y., and E.F. cloned and expressed the recombinant proteins. K.K. and Z.Y. performed the western blotting experiments. K.K., Z.Y., B.B.H., T.M.P.C. and F.Z. labeled antibodies and POIs with dyes or payloads and performed the internalization and cytotoxicity assays. T.M.P.C. performed mass spectrometry to identify DAR of DDCs. K.K.L transduced the Jurkat^PD-1^ line and provided guidance for later experiments. K.K. and J.A.W. wrote the manuscript and all authors reviewed and edited the manuscript.

## Materials and Methods

### KineTAC cloning

Cytokine sequences were sourced from uniprot (uniprot.org). Signal sequences were removed as annotated by uniprot and cloned directly N-terminally of a Trastuzumab hinge, C_H_2 and C_H_3. Kines were either cloned into knob Fc halves with mutations N297G (reduce Fc glycosylation and receptor binding), T366W (knob mutation) or zymeworks B bispecfic Fcs^65^ with LALAPG, T350V T366L, K392L, T394W. Bevacizumab and Cetuximab V_H_ domains^25^ were similarly cloned upstream of a Trastuzumab C_H_1-C_H_3 as a hole Fc half (mutations N297G, T366S, L368A,Y407V) or a zymeworks A Fc half (mutations LALAPG, T366L, K392L, T394W). Initially, an IL-2 signal sequence was used for all constructs. Later, this was switched for the HC5 sequence which greatly enhanced expression yields in most cases.

### Ni IMAC affinity purification

For VEGF^165^ and initial kineTAC expression, the sequence for the VEGF^165^ isoform (A27-R191) was cloned into a pFuse backbone with an N-terminal 10x histidine tag. Plasmids were transiently transfected according to manufacturer specifications Expi293F cells (ThermoFisher scientific) for 6d at 37°C 8% CO_2_ with shaking prior to purification. Cells were pelleted by 4000xg centrifugation and the supernatant collected and 0.2 μm filtered before purifying. Ni Sepharose excel (Cytiva) resin slurry was washed and added to the supernatant at 1:60 final volume. Batch binding was performed with agitation for 1 hr at 4°C with 5 mM imidazole. After flow through, the resin was washed 2x with 6 colume volume (cv) wash buffer (40 mM imidazole in PBS, pH 8.0). Protein was then eluted with 1 cv elution buffer (300 mM imidazole in PBS, pH 8.0) before buffer exchanging into PBS pH 7.4. Standard SDS PAGE gels were used to qualitatively assess purity before proceeding.

### Protein A affinity purification

For later zymeworks based constructs, kineTACs were expressed identically as in Ni IMAC, then purified using HiTrap 1mL prepacked protein A columns (Cytiva). Filtered supernatant was allowed to bind on column at 800 μL/min at rt, washed with PBS at 1mL/min for 8 mins, then eluted with 0.1M Acetic Acid at 500 μL/min for 8 mins. Eluate was then buffer exchanged into PBS pH 7.4 and quantified via A280 and calculated molar absorptivity (ExPasy protparam). Standard SDS PAGE gels were used to qualitatively assess purity before proceeding.

### Cell lines

Cell lines were maintained in T75 (ThermoFisher Scientific) flasks 37 °C and at 5% CO_2_ and split according to ATCC recommendations. FaDu, HeLa, MDA-MB-361, and SiHa were grown in DMEM supplemented with 10% FBS, 1% penicillin/streptomycin, 2mM GlutaMax. Daudi, Daudi^GFP^, and Jurkat^PD-1^ were cultured in RPMI supplemented with 10% FBS, 1% penicillin/streptomycin, 2mM GlutaMax. PBMCs and isolated T and B cells were cultured in complete sRPMI (10% FBS, 1x MEM NEAA, 1 mM sodium pyruvate, 10 mM HEPES, 50 μM 2-mercapto ethanol).

For 96 well plate experiments, only the middle 60 rows were used to avoid edge effects. The surrounding wells were filled with 120 μL PBS. Additionally, the spaces between wells were filled with 100 μL PBS to minimize evaporation.

### PBMC isolation

Human PBMCs were isolated by Ficoll-Paque and maintained in RPMI-1640 with 10% FBS, 1x MEM NEAA, 1 mM sodium pyruvate, 10 mM HEPES, 50 μM 2-mercapto ethanol. Primary CD3+ T cells and CD19+ B cells were isolated from purified PBMCs respectively according to manufacturer protocols using EasySep negative selection kits (Stemcell technologies). Purity was assessed via flow cytometry.

### Protein labeling

VEGF was labeled with pHrodo red succinimyl ester (Sartorius, Gottingen, Germany), pHrodo deep red tetrafluorophenyl ester, or AlexaFluor 647-N-hydroxy succinimide ester (ThermoFisher scientific) following manufacturer recommendations. Briefly, all reactions were carried out at room temperature (rt) in phosphate buffer (100 mM potassium phosphate, 15 mM NaCl, pH 8.0) for 15 mins. For pHrodo red, VEGF conjugates were purified from unreacted dye via spin desalting column (Zeba 7K, ThermoFisher scientific). For pHrodo deep red and AlexaFluor 647, excess dye was simply quenched using equimolar glycine and the crude product used in assays. We found no issues with excess quenched dyes leading to higher background (data not shown).

### ADC conjugation and assays

In vitro ADC assay KineTACs and isotypes were labeled with NHS ester-PEG4-ValCit-PAB-MMAE (BroadPharm, Cat#BP636 25503) with 1:6 molar ratio at RT for 2 h with 100 mM sodium bicarbonate. Antibodies were then desalted using spin desalting columns (Zeba 7K, Thermofisher Scientific). 100,000 Ramos/well were seeded on a 96-well clear plate (Corning, Cat#3917) and ADCs were added and incubated for 72 h. Viability was measured using CellTiter-Glo Reagent 642 (Promega) or via incucyte-based killing assay, where cells were treated with ADCs or DDCs with cytotox green dye (Sartorius, Cat#4633, 1:10000). 0.5% Triton X was added to a control well at 72 hrs to gauge 100% killing.

### ADC characterization

Acetonitrile (HPLC grade), H_2_O (HPLC grade), and formic acid (LC-MS grade) were obtained from Sigma-Aldrich (St. Louis, MO). Liquid chromatography was performed using a Vanquish Flex HPLC system, configured in a direct injection format. Antibodies and antibody drug conjugates (ADCs) were separated on a MAbPac reverse phase polystyrene divinylbenzene HPLC column with 4.0 µm particles and 1,500 Å pore size, with dimensions 1 mm x 100 mm (Thermo Fisher Scientific). During LC separations, mobile phase A (MPA) was 0.2% formic acid (FA) in water and MPB was 80% ACN in water with 0.2% FA. For profiling of the antibodies and ADCs, a 10-min gradient ramped, at a flow rate of 80 µL/min, from 10-70% MPB from 0.1 – 5.1 min, 70-90% MPB from 5.1 – 6.1 min, and held at 90% MPB to 6.9 min before the column was washed and re-equilibrated at 10% MPB for 3 min. During protein separations, the LC column was held at 80 °C. Intact protein MS experiments were performed on a Thermo Scientific^TM^ Orbitrap Eclipse^TM^ Tribrid^TM^ hybrid mass spectrometer system (Orbitrap Eclipse, Thermo Fisher Scientific). Precursors were ionized using electrospray ionization at 2.8 kV with respect to ground. The inlet capillary was held at 350 °C, the vaporizer temperature was 400°C, and the ion funnel RF was held at 60%. Source sheath gas was set to 50 (Arb), auxiliary gas was set to 10 (Arb), and the ion routing multipole was set to 1 mTorr nitrogen gas within intact protein mode. All MS^1^ survey scans were acquired at a resolving power of 15,000 in the Orbitrap analyzer within the high mass range with a scan range of *m/z* 1,000 – 4,000, maximum injection time of 200 ms, and AGC target of 1,000,000 charges using five microscans. In-source fragmentation energy was set to 10 V.For increased signal-to-noise (S/N) ratio in MS spectra, 10-20 individual scans were summed, with averaging performed in the vendor’s postacquisition software (XCalibur FreeStyle, version 1.8). Electrospray MS spectra of all analytes are shown in **Supplemental Fig 3c**,**f**. MS spectra were deconvoluted with XTRACT (Thermo Fisher Scientific) using default parameters and a signal-to-noise ratio threshold of three. Mass spectra were used to identify and integrate peak areas corresponding to the antibody and its MMAE-conjugated species. DAR values were calculated as the weighted average of the number of MMAE drug molecules per antibody based on the relative abundances of each species (**Supplemental Fig 3a-b, d-e**)

### VEGF internalization screen

Fadu and MDA-MB-361 were plated at 17k and 20k cells respectively per well in a 96 well plate. After adhering to the plate for 24 hours, media was removed and 25 nM VEGF-pHrodo red was added simultaneously with 12.5 nM of each kineTAC in 150 μL final volume, one kineTAC per well. An incucyte SX5 (Sartorius, Gottingen, Germany) collected 10x images every 1 hr. Images were analyzed using manufacturer software, and data was plotted as total integrated red intensity over total cell area.

### Degradation experiments

For membrane POIs, cells were plated in 6-or 12-well plates and grown to ∼70% confluency before treatment. Medium was aspirated, and cells were treated with bispecifics or control antibodies in complete growth medium. After incubation at 37 °C for the designated amount of time, cells were washed with PBS, lifted with versene and collected by centrifugation at 300g for 5 min at 4 °C. Samples were then tested by western blotting or flow cytometry to quantify protein levels.

### Flow cytometry

If necessary, cells were lifted using 0.25% Trypsin or versene (versene when surface binding or membrane protein detection was required after lifting; trypsin for elimination of surface bound VEGF). Cells were centrifuged at 1000g for 5 min and washed 3x using flow buffer (PBS, 2% BSA, 0.04% EDTA, pH 7.4). Cells were incubated with primary antibodies or kineTACs diluted in flow buffer for 15 min at 4 °C. Cells were washed three times with cold flow buffer, and secondary antibodies (if applicable) diluted in flow buffer were added and incubated for 30 min at 4 °C. Cells were washed three times with cold flow buffer. Flow cytometry was performed on a CytoFLEX cytometer (Beckman Coulter) using CytExpert software (v2.3.1.22) for data acquisition. Gating was performed on single cells and live cells before acquisition of 10,000 cells. Analysis was performed using FlowJo software (v10.8.0).

### VEGF surface binding removal

Because internalized VEGF-647 is the relevant signal for eTPD, we tested if trypsin could remove confounding surface bound VEGF-647. FaDu cells were incubated with kineTAC and VEGF at 4°C for 1 hour to permit surface binding without internalization, then treated with 0.25% trypsin for 10 min at RT and imaged at 10x once using an incucyte SX5 (Sartorius, Gottingen, Germany). Trypsin removed virtually all surface-bound VEGF-647, validating that post-trypsin signal represents internalized VEGF-647.

### Western blotting

Cell pellets were lysed with 1× RIPA buffer containing cOmplete mini protease inhibitor cocktail (Sigma-Aldrich) at 4 °C for 40 min. Lysates were centrifuged at 16,000g for 10 min at 4 °C, and protein concentrations were normalized by bicinchoninic acid assay (Pierce). Then, 4× NuPAGE LDS sample buffer (Invitrogen) and 2-mercaptoethanol were added to the lysates and boiled for 10 min. Equal amounts of lysates were loaded onto a 4–12% Bis-Tris gel and run at 200 V for 37 min. The gel was incubated in 20% ethanol for 10 min and transferred onto a polyvinylidene difluoride membrane. The membrane was blocked in PBS with 0.1% Tween-20 + 5% BSA for 30min at room temperature with gentle shaking. Membranes were incubated overnight with primary antibodies at respective dilutions at 4 °C with gentle shaking in PBS + 0.2% Tween-20 + 5% BSA. Membranes were washed four times with TBS + 0.1% Tween-20 and co-incubated with horseradish peroxidase goat anti-rabbit IgG (Jackson Immuno Research, 111-035-144; 1:2,000) and 680RD goat anti-mouse IgG (LI-COR, 926-68070; 1:10,000) in PBS + 0.2% Tween-20 + 5% BSA for 1 h at room temperature. Membranes were washed four times with TBS + 0.1% Tween-20 and then washed with PBS. Membranes were imaged using an OdysseyCLxImager (LI-COR). SuperSignal West Pico Plus chemiluminescent substrate (ThermoFisher Scientific) was then added and imaged using a ChemiDoc Imager (Bio-Rad). Band intensities were quantified using Image Studio software (LI-COR, v5.2.5).

### PBMC assay

PBMCs were treated at 200k cells/well in 96 well plates in sRPMI with indicated kineTACs or isotypes with or without VEGF-AlexaFluor647 at 25 nM. For surface binding experiments, no VEGF was added and dosing was done at 4 °C for 15 min. For internalization experiments, kineTACs and VEGF were simultaneously added to cells and allowed to incubate at 37 °C for 24 hours before analyzing by flow cytometry. All cells were gated as single live cells, then T cells were gated as CD3/CD5 positive and CD19 negative, and B cells were gated as CD19 positive and CD3/CD5 negative. Percent positivity was defined by approx 1% positive in the VEGF or isotype only condition and applied as a gate to all samples of that type.

### T cell assays

T cells were isolated from frozen PBMC using EasySep Human T Cell Isolation kit (Stemcell Technologies) and subcultured with 30U/mL IL2 in sRPMI. T cells were first activated on plates coated with 1 µg/mL OKT3 and anti-human CD28 (BioLegend) in the presence of 30U/mL IL2. After 1 day, kineTACs were added to induce the degradation of PD-1 and incubated for another 24 hours. At the end of the treatment cells were harvested for western blot analysis.

### Heatmap creation

Tissue expression profiles were pulled from the Human Protein Atlas^23^ IHC data, based on staining in 76 cell types for 45 normal tissues. Numbers were assigned to each qualitative expression level (None:0, Low:1, Medium:2, High:3) for the purposes of averaging across cell types in each tissue. Expression levels were then displayed for relevant cytokine receptors in a heatmap in python using matplotlib.

### PI and purity analyses

PI was calculated using amino acid sequences of the full kineTACs in ExPasy Protparam (https://web.expasy.org/protparam/). Purity was estimated from SDS PAGE gels. Total yield was estimated from A280 and volume measurements.

#### Antibodies used included

rabbit anti-human EGFR (Cell Signaling Technology, Cat# 4267S, 1:1000), rabbit anti-human PD-1 (Cell Signaling Technology, Cat# D4W2J, 1:1000), mouse anti-human IL7R (R&D Systems, Cat# MAB306, 1:1000), rabbit anti-human phospho NF-κB p65 Ser 536 (Cell Signaling Technology, Cat#93H1, 1:1000), mouse anti-human NF-κB p65 (Cell Signaling Technology, Cat#93H1, 1:1000), mouse anti-human β-actin (Cell Signaling Technology, Cat# 8H10D10, 1:1000), and mouse anti-human β-tubulin (Cell Signaling Technology, Cat# DM1a, 1:1000). Secondary antibodies for western blot included IRDye 800CW goat anti-rabbit IgG (LI-COR Biosciences, Cat# 925-32210, 1:1000), IRDye 680RD goat anti-mouse IgG (LI-COR Biosciences, Cat# 925-68070, 1:10000), peroxidase-conjugated goat anti-rabbit IgG (H+L) (Jackson ImmunoResearch, Cat# 111-035-144, 1:2000), and peroxidase-conjugated goat anti-mouse IgG (H+L) (Jackson ImmunoResearch, Cat# 115-035-003, 1:2000).

#### Flow cytometry antibodies used included

FITC mouse anti-human CD3 (BioLegend, Cat# 300405), FITC mouse anti-human CD5 (BioLegend, Cat# 364021), PE mouse anti-human CD14 (BioLegend, Cat# 301805), BV421 mouse anti-human CD19 (BioLegend, Cat# 302233), PE mouse anti-human CD33 (BioLegend, Cat# 303403), APC mouse anti-human IL4Rα (BioLegend, Cat# 355005), PE mouse anti-human IL4Rα (BioLegend, Cat# 355003), APC mouse anti-human IL7Rα (BioLegend, Cat# 351315), FITC mouse anti-human IL7Rα (BioLegend, Cat# 351311), PE mouse IgG1 isotype control (BioLegend, Cat# 400211), FITC mouse IgG1 isotype control (BioLegend, Cat# 400107), and Alexa Fluor 647-conjugated Protein A (ThermoFisher Scientific, Cat# P21462).

