## Supplemental Data for "A cytokine receptor-targeting chimera (kineTAC) toolbox for expanding extracellular targeted protein degradation"

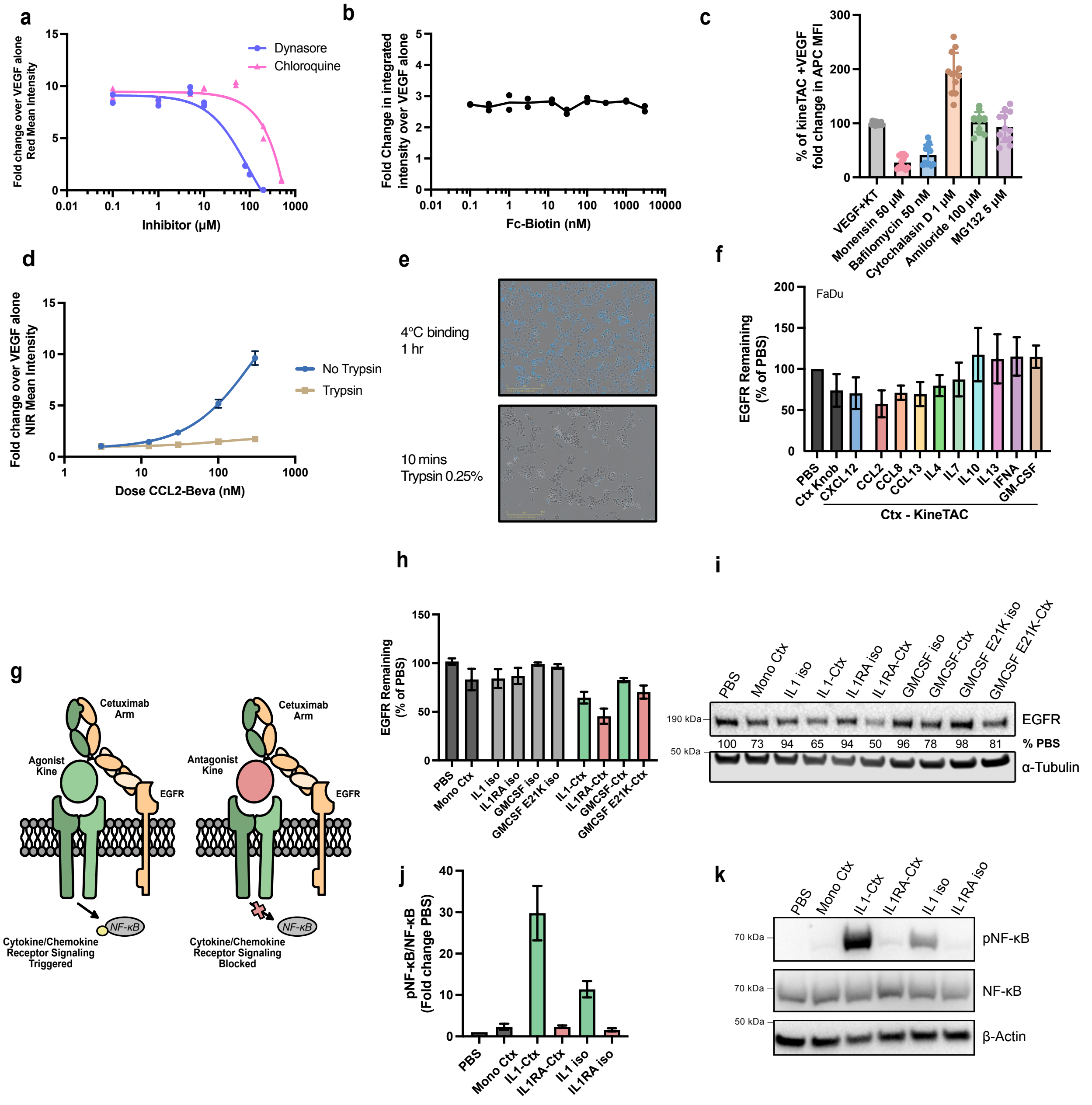


**Supplemental Fig 1 | Additional screen and assay validation and antagonism follow up. a,** Incucyte analysis for treating cells with 25 nM of VEGF-pHrodo red and 25 nM of IL7-Bevacizumab kineTAC and a dose response of dynasore or chloroquine at 24 hr. Data are from two biological replicates. **b**, Incucyte analysis of VEGF-pHrodo red 25 nM + IL7-Bevacizumab kineTAC 25 nM with a dose response of Fc-biotin (no Fab) at 24h. Data represents two biological replicates. **c**, flow cytometry data of 100nM kineTAC + 20nM VEGF-647 24 hr internalization with concurrent inhibitor (name and concentration displayed) treatment in FaDu cells. Data is plotted as a percentage of uptake triggered by VEGF and kineTAC (VEGF+ KT) without pharmacological inhibitor (i.e, reduction in kineTAC uptake with inhibitor). Data is an aggregate of two technical replicates for CCL2, CCL8, CXCL12, IL4, IL7, and IL21 Bevacizumab kineTACs. Error bars represent SEM. **d-e**, Incucyte analysis of CCL2-Bevacizumab dose response with 12.5 nM VEGF-647 after 4°C binding for 1h. For trypsin treated cells, each well was incubated with 0.25% trypsin for 10 mins and quenched with complete DMEM. For the no trypsin treatment, cells were simply washed with PBS and imaged. **e,** representative images from part **d** with scale bars shown. Blue = NIR (VEGF-647) signal. **f**, Western blot quantification of mean β-actin normalized EGFR levels as a percent of PBS only treatment after 24 hr with 50 nM kineTAC or isotype treatment in FaDu cells. Ctx Knob = monomeric cetuximab (Cetuximab Half IgG Hole with Fc only Knob IgG). Error bars are SEM. **g**, schematic of antagonist and agonist kineTACs targeting EGFR. In one case, an agonist cytokine presumably triggers signaling of the interacting cytokine/chemokine receptor. In the antagonist case, the opposite is true. **h-i,** quantification (**h**) and representative western blot (**i**) and  of EGFR levels in HeLa cells after 24 hr treatment with 100 nM of the indicated kineTAC. Data is normalized to α-tubulin from three biological replicates and corresponding SEM is displayed. **j-k**, Quantification (**j**) and representative western blot (**k**) of phospho NF-κB (p65, Ser536), NF-κB, and β-Actin levels in HeLa cells after 15 min treatment with 100 nM of the indicated kineTAC. Data is normalized to β-actin from three biological replicates and corresponding SEM is displayed.

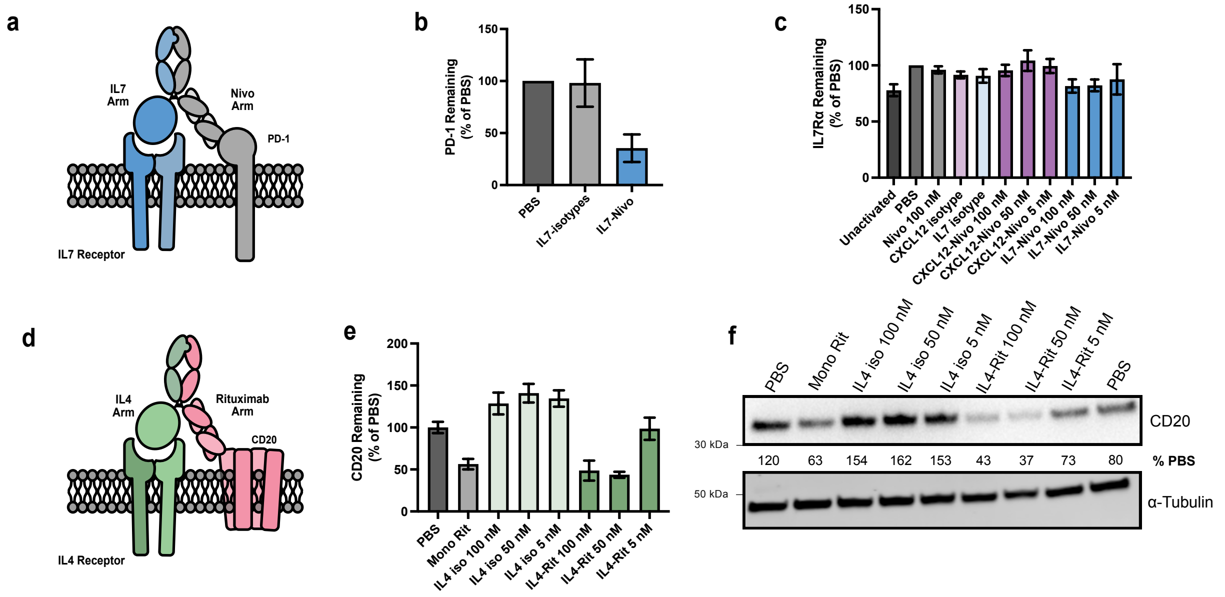


**Supplemental Fig 2 | Further kineTAC applications. a**, Schematic of IL7-Nivolumab (Nivo) kineTAC. IL7 triggers the internalization and degradation of PD-1 through the IL7 receptor. **b**, Quantification of western blot PD-1 levels from **Fig 3d.** Jurkat PD-1 overexpression cell line after 24 hr with 50 nM of the displayed kineTAC or isotype antibody control. The PD-1 signal was normalized to β-Actin as a percentage of the PBS only normalized PD-1 signal is displayed. PD-1 signal is HRP luminescence and β-actin is from a LICOR secondary. IL7 isotypes = half IL7 knob IgG with Fc only hole IgG, or free IL7. Data from two biological replicates. **c**, Western blot quantification of activated primary T cell mean β-actin normalized IL7Rα levels as a percentage of the PBS only control after 24 hr treatment with kineTAC at displayed concentrations or isotypes at 100 nM. Five total donors represented, from the same blots as **Fig 3f. d**, Schematic of IL4-Rituximab (Rit) kineTAC. IL4 triggers the internalization and degradation of CD20 through the IL4 receptor. **e-f,** quantification (**e**) and representative western blot (**f**) of CD20 levels in Ramos cells after 24 hr treatment with of the indicated kineTAC at indicated concentrations. CD20 %PBS is normalized CD20 levels for given treatment as a percentage of average normalized PBS CD20 signal per blot. Mono Rit = monomeric Rit, containing Rit Half IgG Hole with Fc only Knob IgG, at 100 nM. Data is normalized to α-tubulin from three biological replicates. All error bars represent SEM.

**a**

| # MMAE | z | m/z | Intensity | Relative Intensity (%) |
|---|---|---|---|---|
| 0 | 42 | 1746.54 | 1.51E+06 | 52.95 |
| 1 | 42 | 1779.42 | 8.00E+05 | 28.05 |
| 2 | 42 | 1812.31 | 3.62E+05 | 12.69 |
| 3 | 42 | 1845.03 | 1.80E+05 | 6.31 |
| SUM | | | 2.85E+06 | |
| | | | DAR | 0.72 |

**b**

| # MMAE | z | m/z | Intensity | Relative Intensity (%) |
|---|---|---|---|---|
| 0 | 41 | 1789.14 | 9.64E+05 | 33.46 |
| 1 | 41 | 1822.79 | 1.26E+06 | 43.73 |
| 2 | 41 | 1856.49 | 5.12E+05 | 17.77 |
| 3 | 41 | 1889.54 | 1.45E+05 | 5.03 |
| SUM | | | 2.88E+06 | |
| | | | DAR | 0.94 |

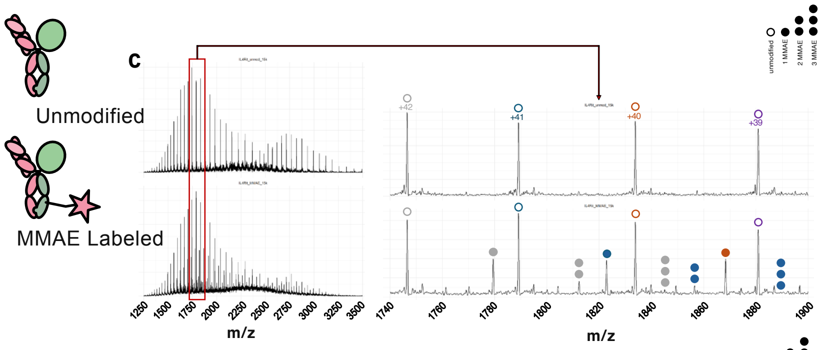


**d**

| # MMAE | z | m/z | Intensity | Relative Intensity (%) |
|---|---|---|---|---|
| 0 | 51 | 2024.28 | 3.04E+06 | 34.86 |
| 1 | 51 | 2051.55 | 2.47E+06 | 28.33 |
| 2 | 51 | 2078.61 | 2.00E+06 | 22.94 |
| 3 | 51 | 2105.81 | 1.21E+06 | 13.88 |
| SUM | | | 8.72E+06 | |
| | | | DAR | 1.16 |

**e**

| # MMAE | z | m/z | Intensity | Relative Intensity (%) |
|---|---|---|---|---|
| 0 | 50 | 2064.77 | 3.24E+06 | 33.68 |
| 1 | 50 | 2092.53 | 2.68E+06 | 27.86 |
| 2 | 50 | 2120.06 | 1.91E+06 | 19.85 |
| 3 | 50 | 2147.79 | 1.79E+06 | 18.61 |
| SUM | | | 9.62E+06 | |
| | | | DAR | 1.23 |

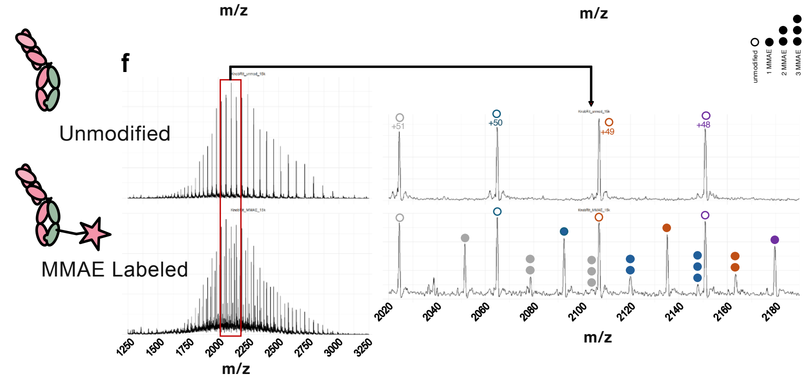


**Supplemental Fig 3 | Characterization of IL4-Rituximab DDC. a-b,** IL4-Rituximab z, m/z, integrated ion intensities, and percentage of total intensity for a given number of MMAE labeling events. Drug to antibody ratio (DAR) reported in the bottom right. Separated into two tables, one for each of the top two z values. **c,** full mass spectrum (left) and detailed range of interest highlighted in red (right) for unmodified (top) and MMAE labeled (bottom) IL4-Rituximab. Dots indicate number of labeling events by calculated mass. Colored by charge state (+42 gray, +41 blue, +40 orange, +39 purple). **d-e,** similar to (**a-b**), but for monomeric rituximab isotype. **f,** similar to (**c**) but for monomeric rituximab isotype. Colored by charge state (+51 gray, +50 blue, +49 orange, +48 purple).

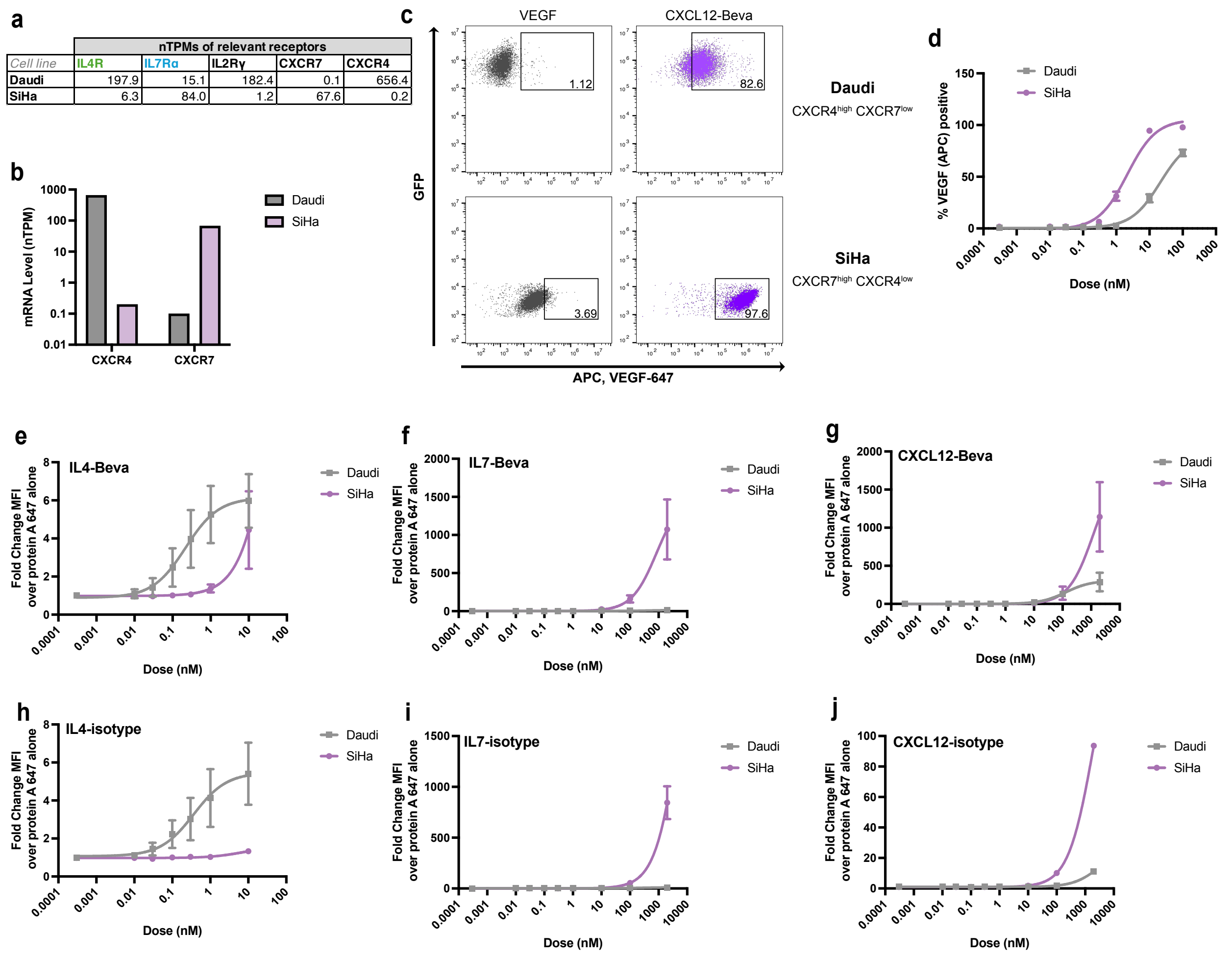


**Supplemental Fig 4 | Further analysis of cell specific uptake. a,** Table showing normalized RNA TPMs extracted from Human Protein Atlas for relevant cytokine receptors in the Daudi and SiHa cells. **b,** normalized TPMs for the two CXCL12 receptors, CXCR4 and CXCR7 in Daudi and SiHa cells from Human Protein Atlas. **c,** Representative flow cytometry data from coculture experiment. Approximately equal numbers of Daudi and SiHa cells were incubated for 24 hrs with 25 nM VEGF-647, and 10 nM CXCL12-Bevacizumab. Gates show thresholds for VEGF positivity, defined as about 1% for VEGF-647 only. Top row: FITC $^{+}$ Daudi$^{GFP}$. Bottom row: FITC $^{-}$ SiHa. Percentages in gate are displayed. **d,** Coculture VEGF-647 percent positivity (same gate as in **c**) as a dose response of CXCL12-Bevacizumab**.** Mean values and corresponding ± SEM from three biological replicates. Curves are three parameter non-linear regression best fits. **e-j,** 4°C surface binding experiments in monocultures of Daudi or SiHa cells that prevents internalization. Indicated kineTAC or isotype was dosed for 15 min at 4°C, then washed and stained by protein A-AlexaFluor647. Fold change in median fluorescence intensities of protein A binding above protein A-AlexaFluor647 alone is presented, indicating surface bound kineTAC.

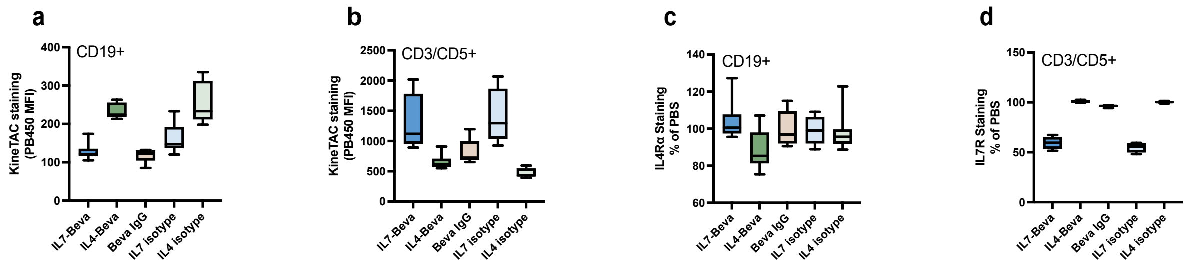


**Supplemental Fig 5 | Surface binding and receptor blocking of kineTACs in PBMCs. a-b,** PBMCs from 4 donors were incubated with 10 nM indicated kineTAC at 4°C for 15 min, then washed and stained with protein A-BV421 (**a-b**), α-IL7R (**c**), or α-IL4R (**d**). **a,b** box plots of the median fluorescence intensity (MFI) of protein A-BV421, representing kineTAC binding and **c,d** box plots of the reduction in α-IL7R MFI or α-IL4Rα MFI after treatment with the indicated kineTAC as a percentage of the PBS MFI. **a** and **c** are from the $CD19^+$, $CD3/CD5^-$ gates and **b** and **c** are from the $CD19^-$ $CD3/CD5^+$ gates (same gates as in **Figure 5g-h**). Box plots depict minimum, 1st quartile, median, 3rd quartile, and maximum of from three technical replicates of four different donors.

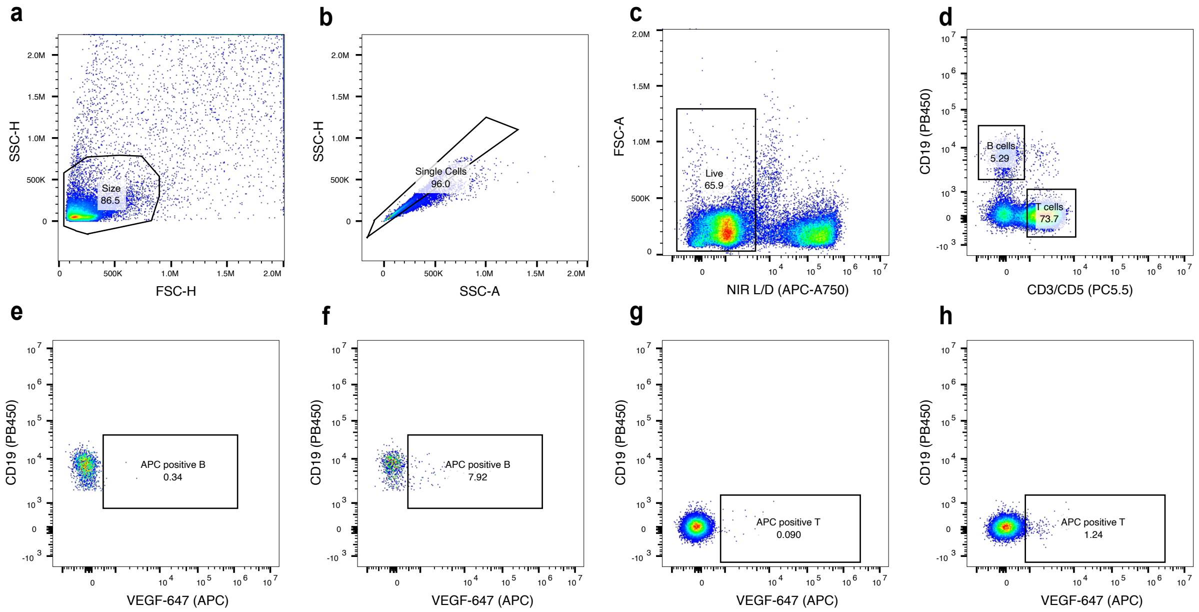


**Supplemental Fig 6 | Gating strategy for PBMC experiment.** Gates from **Figure 5g-h** are shown for a representative donor. **a,** cell size gate. **b**, singlet gate. **c**, live dead gate based on near infra red (NIR) live/dead (L/D) amine reactive dye. **d**, B and T cell gates, based on CD3/CD5 staining (x axis) and CD19 staining (y axis). **e**, VEGF-647 positive gate definition for B cells, based on the vehicle treated (no VEGF) control. **f**, same as (**e**), but with 25 nM VEGF-647 added. **g**, VEGF-647 positive gate definition for T cells, based on the vehicle treated (no VEGF) control. **h**, same as (**g**), but with 25 nM VEGF-647 added.

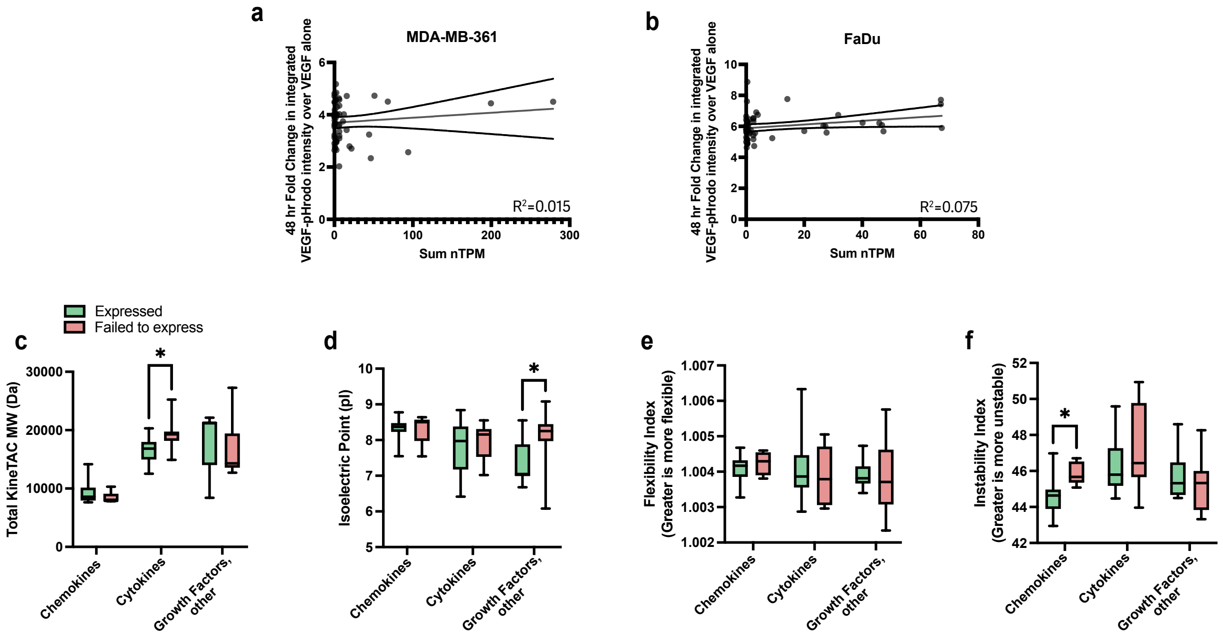


**Supplemental Fig 7 | Expression and screen correlations. a-b,** correlations between the sum of mRNA transcript levels of all known receptors for a given kineTAC from Human Protein Atlas[1] and the 48 hour fold change uptake of VEGF-pHrodo (from **Fig 2b**) in MDA-MB-361 (**a**) and in FaDu (**b**). **c-f**, biophysical parameters of both expressed (used in screen, see **Fig 1c**) and non-expressed (excluded from screen) cytokines, chemokines, and growth factors. Total KineTAC molecular weight (**c**), calculated isolectric point (**d**), Flexibility Index (**e**) and instability index (**f**) calculated using biopython ProtParam module. Asterisks represent $p < 0.05$ from unpaired t test. All other comparisons within families are non-significant.
